# The classical MHC class I and II genes of *O. m. formosanus* exhibit different polymorphism levels

**DOI:** 10.1101/2025.03.31.646510

**Authors:** Zhi-Wei Zhang, Lukas A. Fugmann, Shu Yuan Yang, Jin-Chywan Gwo, Sebastian D. Fugmann

## Abstract

*Oncorhynchus masou formosanus* (Formosa landlocked salmon) is a salmonid fish endemic to Taiwan with a critically endangered extant small population in high-altitude mountain streams. To begin to assess the remaining genetic diversity we characterized the single classical MHC class I (*UBA*) and MHC class II (*DAA* and *DAB*) genes of teleost fish in a small cohort of eight unrelated *O. m. formosanus* individuals. We focused on the exons encoding the peptide binding regions of these complexes as they are considered the most highly polymorphic regions in vertebrate genomes. Surprisingly, the *DAA* and *DAB* genes appeared homozygous and identical among all eight fish indicating that the encoded class II complex is monomorphic. In contrast, three distinct *UBA* alleles were discovered with one dominant allele present in every single individual. Notably, 75% were heterozygous indicating a selective advantage of heterozygosity at this locus. Lastly, our MHC alleles differ from those present in the genome of the closely related Japanese *O. m. masou*, suggesting that the loss of DAA/DAB diversity and the emergence of the dominant UBA allele occurred after their populations were isolated. Together we discovered residual genetic diversity at the classical MHC class I locus in *O. m. formosanus* and maintaining this allelic variation unlike the homozygous *DAA* and *DAB* genes is likely important for its survival in its geographically restricted unique habitat.

## 1. Introduction

*Oncorhynchus masou formosanus* (Formosa landlocked salmon) is a salmonid fish that is endemic to Taiwan where it resides exclusively in the Chichiawan tributary that is part of the upper reaches of the Tachia river in the central mountains of Taiwan. This single isolated population shares common ancestors with the *Oncorhynchus masou* species in the rivers and lakes of Japan and neighboring countries bordering the North Pacific. Their split occurred after the last glacial period with the increasing water temperatures in the lower reaches of the Tachia river trapping an *O. masou* population in the cold water of high-altitude mountain streams (Healey et al., 2011). Hence *O. m. formosanus*, just like the landlocked *O. m. “Biwa”* (Biwa trout) in Lake Biwa in Japan, adopted a non-anadromous lifestyle while most other *O. masou* populations remain anadromous (Morita, 2011). The continuous destruction of its native habitat in the 20^th^ century resulted in a collapse in the total number of *O. m. formosanus* down to as low as a few hundred and thus created a second genetic bottleneck. As recent conservation efforts for this critically endangered fish including breeding in a hatchery helped to boost the population levels significantly, it now raises the question about how much genetic diversity remains in the extant population in the wild (Gwo et al., 2008; Yamamoto et al., 2020).

The genes encoding the components of the major histocompatibility complexes (MHCs) show the highest level of polymorphisms of all genes in vertebrate genomes (Jin et al., 2018), and they are thus frequently used to study population (Sommer, 2005). MHC protein complexes are expressed on the surface of antigen-presenting cells to present pathogen peptides to T lymphocytes during immune responses. While the peptide-binding region (PBR) of classical MHC class I complexes is formed by a single heavy chain gene that binds to proteasome derived peptides from the intracellular space, the PBR of classical MHC class II complexes formed by an αβ heterodimer that present peptides from the extracellular space that are generated in phagolysosomes. Importantly, the sequence variations of the MHC complexes largely occur in the α-helices that line the PBR, contact the pathogen peptides and confer peptide specificity for each MHC allele (Stern and Wiley, 1994). The polymorphic nature of both MHC complexes is beneficial for population level immunity and hence survival (Apanius et al., 1997). There are, however, also closely related non-classical MHCs that do not present peptides and those or typically monomorphic or oligomorphic.

In most salmonid fish there is only a single classical MHC class I complex (with its heavy chain encoded by the *UBA* gene) and a single classical MHC class II complex (encoded by the linked *DAA* and *DAB* gene pair) (Grimholt and Lukacs, 2021). All three genes are polymorphic in all *Oncorhynchus* species that have been analyzed with UBA allele numbers of up to 34 in *Oncorhynchus nerka* (sockeye salmon) (McClelland et al., 2011) and ten or more alleles of DAA and DAB in *Oncorhynchus mykiss* (rainbow trout) (Gómez et al., 2010). But no such information is available for *O. m. formosanus* or any of the Japanese *O. masou* as of yet. We thus decided to investigate these classical MHC genes in a cohort of *O. m. formosanus* individuals collected in the wild in 2016 (and not the potentially less diverse hatchery stock). As no genome assembly exists for this species, we first cloned and sequenced the relevant exons of the *OmfoDAA*, *OmfoDAB*, and *OmfoUBA* genes. The subsequent analysis of their diversity in a small sample set of eight fish indicated that both MHC class II genes were monomorphic but distinct from the genes in the recent first Japanese *O. m. masou* genome assembly (Christensen et al., 2025). In contrast, we discovered a total of three alleles of the *OmfoUBA* locus in our cohort with an observed heterozygosity of 0.750.

## 2. Material and Methods

### 2.1 Oligonucleotides

All oligonucleotides (Suppl. Table 1) were synthesized by a commercial provider (Genomics, Taiwan).

### 2.2 Genomic DNA

*O. m. formosanus* is an endangered species and to minimize disturbance to this species in the wild we executed this study using pre-existing ethanol preserved tissue samples (obtained by Prof. JC Gwo, National Taiwan Ocean University). Ethanol preserved fin tissue sample from a eight individuals (FLS#775 [male, body length 270 mm, weight 205 g], FLS#776 [male, 256 mm, 160 g], FLS#810 [male, 360 mm, 535 g], FLS#815 [male, 263 mm, 220 g], FLS#842 [male, 283 mm, 260 g], FLS#844[male, 280 mm, 200 g], FLS#848 [male, 305 mm, 290 g], FLS#850 [male, 280 mm, 235 g]) was minced with a scalpel, and genomic DNA was purified using the EasyPure Genomic DNA Spin Kit (Bioman).

### 2.3 PCR

PCR reactions were conducted with Phusion polymerase (ThermoScientific) using the primer pairs listed in Suppl. Table 1, and 100 ng genomic DNA as the template. The PCR programs to amplify the *OmfoDAA and OmfoDAB* genomic fragments started with an initial denaturation step at 98°C for 2 min followed by 35 cycles of 98°C for 20 sec, annealing at 54°C for 20 sec, and extension at 72°C for 1 min; a final extension step at 72°C for 7 min concluded the reaction. For the amplification of *UBA* exon 2 and exon 3+4 fragments the extension time was shortened to 30 sec. To avoid PCR artefacts, for all individual genomic DNA samples two separate PCR reactions were conducted per primer pair.

### 2.4 Cloning and sequencing of PCR products

The PCR products were resolved on 1% agarose gels and the bands for each fish/gene were excised and pooled prior to purification using the EasyPure PCR/Gel Extraction Kit (Bioman). Purified fragments were ligated into pJET1.2 vector using the CloneJET PCR Cloning Kit (ThermoScientific) and the ligation reactions were transformed into DH5α *E. coli*. A minimum of eight colonies was picked for each cloned amplicon and used to inoculate 2 ml liquid cultures in LB broth with 100 µg/ml ampicillin. Plasmids were purified from overnight cultures with the Tools Plasmid Mini Kit (Biotools), and positive clones were identified by restriction digests. The sequences of the cloned PCR products were determined by a commercial Sanger sequencing service (Genomics, Taiwan).

### 2.5 Bioinformatics

The pair-wise sequence comparisons of the *O. m. formosanus MHC* genes with the respective genomic loci from other *Oncorhynchus* species and of the corresponding amino acid sequences were conducted using the BLAST server at NCBI (https://blast.ncbi.nlm.nih.gov/Blast.cgi). Multiple sequence alignments were conducted using CLUSTALW (Larkin et al., 2007) with default parameters at the webserver. The amino acid sequence of the DAA, DAB, and UBA alleles are listed in Supplementary Data file 1.

### 2.6 Sequence information

All annotated gene sequences and the respective transcript and protein sequences were obtained from NCBI Genbank (ncbi.nlm.nih.gov) and Ensembl (www.ensembl.org). The Genbank accession numbers for the *O. m. formosanus* DAA, DAB, UBA exon 2, and UBA exon3+4 fragments cloned and sequences in this study are XXXX (*OmfoDAA*), XXXX (*OmfoDAB*), XXXX (*OmfoUBAα1*01:01*), XXXX (*OmfoUBAα1*02:01*), XXXX (*OmfoUBAα1*03:01*), XXXX (*OmfoUBAα2*01:01*), XXXXX (*OmfoUBAα2*02:01*), XXXX (*OmfoUBAα2*03:01*).

## 3. Results

### 3.1 The DAA and DAB genes of O. m. formosanus are monomorphic

The first step towards assessing the MHC diversity in *O. m. formosanus* was to design PCR primer pairs that would allow amplifying the exons *DAA* and *DAB* genes that encode the PBRs (the α1 domain of DAA, and the β1 domain of DAB, respectively) of the classical MHC class II complexes (Fig. 1). As we relied on genomic DNA samples and wanted to obtain as much protein coding sequence as possible, we selected the primers in the intronic regions flanking *DAA* exons 2 and 3, and *DAB* exons 2 and 3 based on sequence conservation across three Pacific salmon species (*O. mykiss*, *O. nerka*, and *Oncorhynchus kisutch [coho salmon]*). We started with the genomic DNA of one individual (FLS#775) as a template and readily obtained PCR products of the expected sizes for both primer pairs. To confirm the identity of these DNA fragments and to obtain the relevant coding sequences, each PCR product was ligated into a cloning vector and the nucleotide sequences of at least six clones for each amplicon was determined.

**Figure 1.**
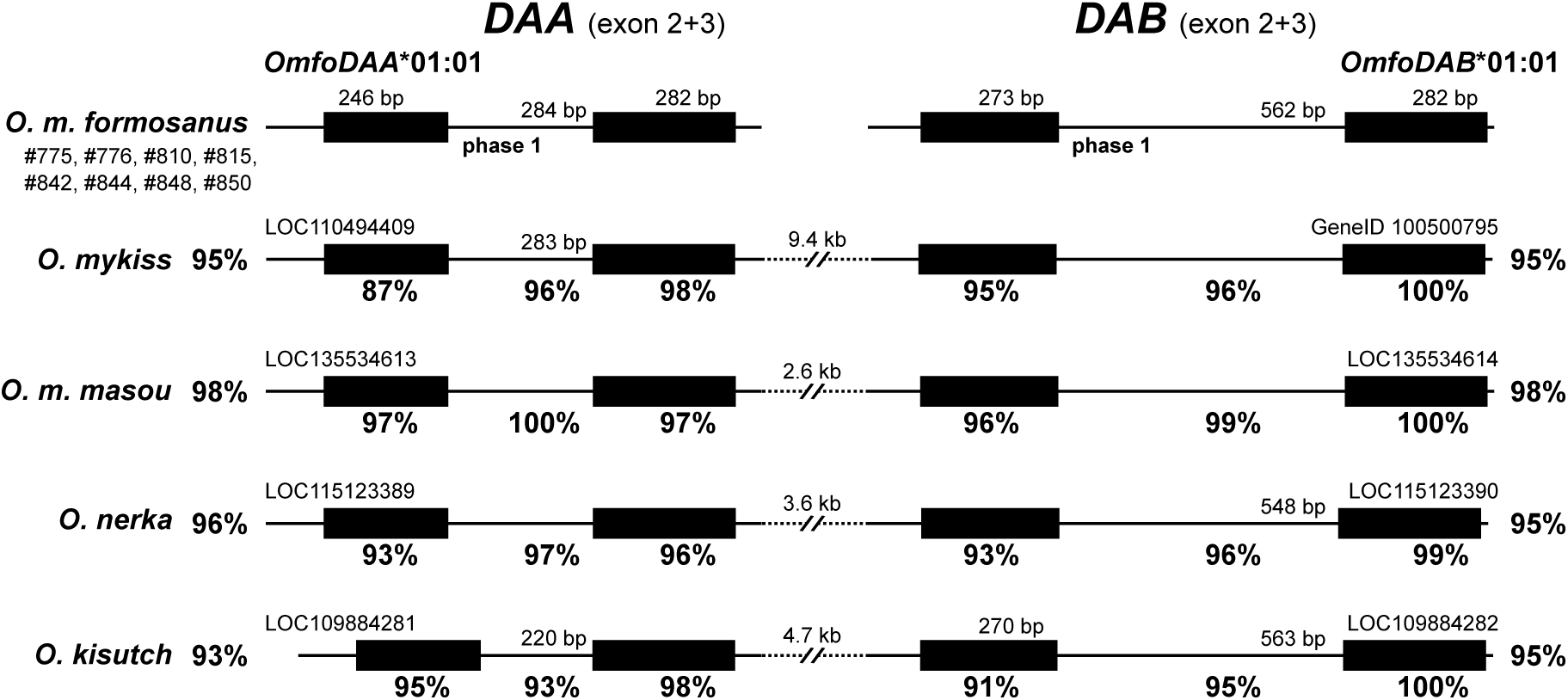
Comparison of the DAA and DAB loci of O. m. formosanus to those of other Pacific salmon species. The PCR amplicons of exons 2 and 3 of *OmfoDAA* and *OmfoDAB* are compared to the respective regions in the genome assemblies of *O. mykiss* (USDA_OmykA_1.1, GCF_013265735.2), *O. m. masou* (UVic_Omas_1.1, GCF_036934945.1), *O. nerka* (Oner_Uvic_2.0, GCF_034236695.1), and *O. kisutch* (Okis_V2, GCF_002021735.2). The overall level of sequence identity to the *O. m. formosanus* PCR products (which were identical for all the individuals analyzed) compared to the respective alleles present in the genome assemblies in the indicated species is shown in front (*DAA*) or after (*DAB*) each gene locus (note that a manual annotation of *OmykDAB* is shown here as this gene model in USDA_OmykA_1.1 lacks exon 1 and 2 and is mis-named as *oncmyk-dbb*). The sizes of the exons and introns are shown above and the percentages of sequence identity below each element. All sizes identical to those observed in *O. m. formosanus* have been omitted for clarity.

Interestingly, the six sequences for each of the class II genes (*DAA* and *DAB*) were identical suggesting that this individual fish was homozygous for *DAA* and *DAB*. To assess the level of allelic variants in *DAA* and *DAB* within the *O. m. formosanus* population, the genomic DNA from ethanol-preserved fin tissues of seven additional individuals were used as templates to amplify the respective MHC gene fragments. To obtain the accurate nucleotide sequence of both alleles (from heterozygous individual) all PCR amplicons were again cloned and multiple clones from each fish were sequenced. Surprisingly, the DAA and DAB sequences of all fish were all identical to those of FLS#775 suggesting that at least within our sample size of eight there is only a single allele (hereafter referred to as OmfoDAA*01:01 and OmfoDAB*01:01, respectively) that all individuals are likely homozygous. Although it remains formally possible that our primers were unable to pick up the second allele of *DAA* and *DAB* in every single fish, it is worth noting that all four primer binding sites are highly conserved across three *Oncorhynchus* species and hence it is unlikely that our reagents would fail amplify to any other alleles. To test whether we in fact amplified DAA and DAB fragments and not any of the non-classical MHC class II genes that tend to be monomorphic, we conducted an unbiased BLAST search in the genomes of the three Pacific salmon species (*O. mykiss*: USDA_OmykA_1.1, *O. nerka*: Oner_Uvic_2.0, *O. kisutch*: Okis_V2) based on which we designed the primers and the *O. m. masou* genome (UVic_Omas_1.1) that became recently available. Both amplicons were highly similar to their *DAA* and *DAB* genes with identities ranging from 93-98% and 95-98%, respectively (Fig. 1). Notably there are four unplaced genomic scaffolds for the *O. m. masou* genome assembly that contain *DAA*- and *DAB*-like gene loci, but both genes are only functional in one of them (scaffold_3666, NW_027010068.1) which we consider to be *OmmaDAA* (LOC135534614) and *OmmaDAB* (LOC135534614) hereafter. In contrast, scaffold 1763 (NW_027007802.1) harbors a closely related gene pair (LOC135532195 and LOC135532194, respectively) with the latter carrying a frame shift mutation in exon 2. In addition, contigs 9151 (NW_027015628.1) and 6349 (NW_027012780.1) appear to be unlinked fragments of the same chromosomal region lacking exon 1 of their DAA-like gene (LOC135538130) exhibiting multiple point mutations within exons 2 and 3 of their DAB-like pseudogene (LOC135536568). It is possible that some (if not all) of these genes are assembly artefacts where the genome assembly algorithm did not coalesce the maternal and paternal allele into a single haploid chromosomal region.

A systematic comparison and annotation of our sequences to the *DAA* and *DAB* loci from four *Oncorhynchus* species revealed an almost perfect conservation of exon and intron length and a perfect conservation of the exon-intron boundaries and intron phasing (Fig. 1). The exons 3, in particular of *DAB*, encoding part of the Ig domain at the base of the extracellular domain of each MHC class II subunit are more conserved compared to the exon 2 sequences that allow for altered peptide binding specificity for each allele. As we only included the *DAA* and *DAB* alleles present in the respective haploid genome sequence assemblies, the similarities to other alleles of exon 2 could be even higher or lower.

To extend this analysis to the amino acid sequence level, we retrieved, linked, and translated the exons sequences within our *DAA* and *DAB* amplicons. As expected, a BLASTP comparison with the comprehensive *O. mykiss* MHC sequence datasets at GenBank and in the IPT-MHC database revealed high similarity to the DAA (up to 87% identity in case of OnmyDAA*03:01) and DAB (up to 94% in case of OnmyDAB*08:01) sequences but much lower similarity to the non-classical MHC class II subunits DBA (45%, XP_036793547.1), and DBB (49%, XP_021479592.1) in *O. mykiss*. This indicates that we cloned the *OmfoDAA* and *OmfoDAB* sequences and not any distantly related non-classical (and likely monomorphic) MHC class II genes. Note that a comparison with DBA and DBB from *O. m. masou* was not possible as no orthologs of these proteins are encoded in the current genome assembly of that fish. Comparing our *O. m. formosanus* sequences to all salmonid fish MHC sequences at GenBank revealed that in line with the phylogenetic relationship of these species the closest best matches were the *O. m. masou* DAA (94%, XP_064817619.1), and DAB (95%, XP_064817620.1), respectively. While only one of the five differences in the DAA PBR is at a position predicted to be directly contacting the antigen peptide (Fig. 2A), this ratio is much higher (4/9) for the DAB PBR (Fig. 2B). Similarly, the alignment with the most similar *O. mykiss* alleles revealed that the majority of differences located to the PBR (DAA: 17/21, DAB: 10/11) with 7/15 predicted peptide contact residues of DAA being affected, but only 1/12 for DAB (Fig. 2). Within PBRs the sequence variations are relatively evenly distributed across the β-sheet (DAA:8/17 DAB: 6/17) forming its base and the α-helices flanking the antigen peptide (DAA:9/17, DAB 4/17). As expected, the sequences of the Ig domain encoded by exon 3 are much more conserved with only four (DAA) and one (DAB) differences.

**Figure 2.**
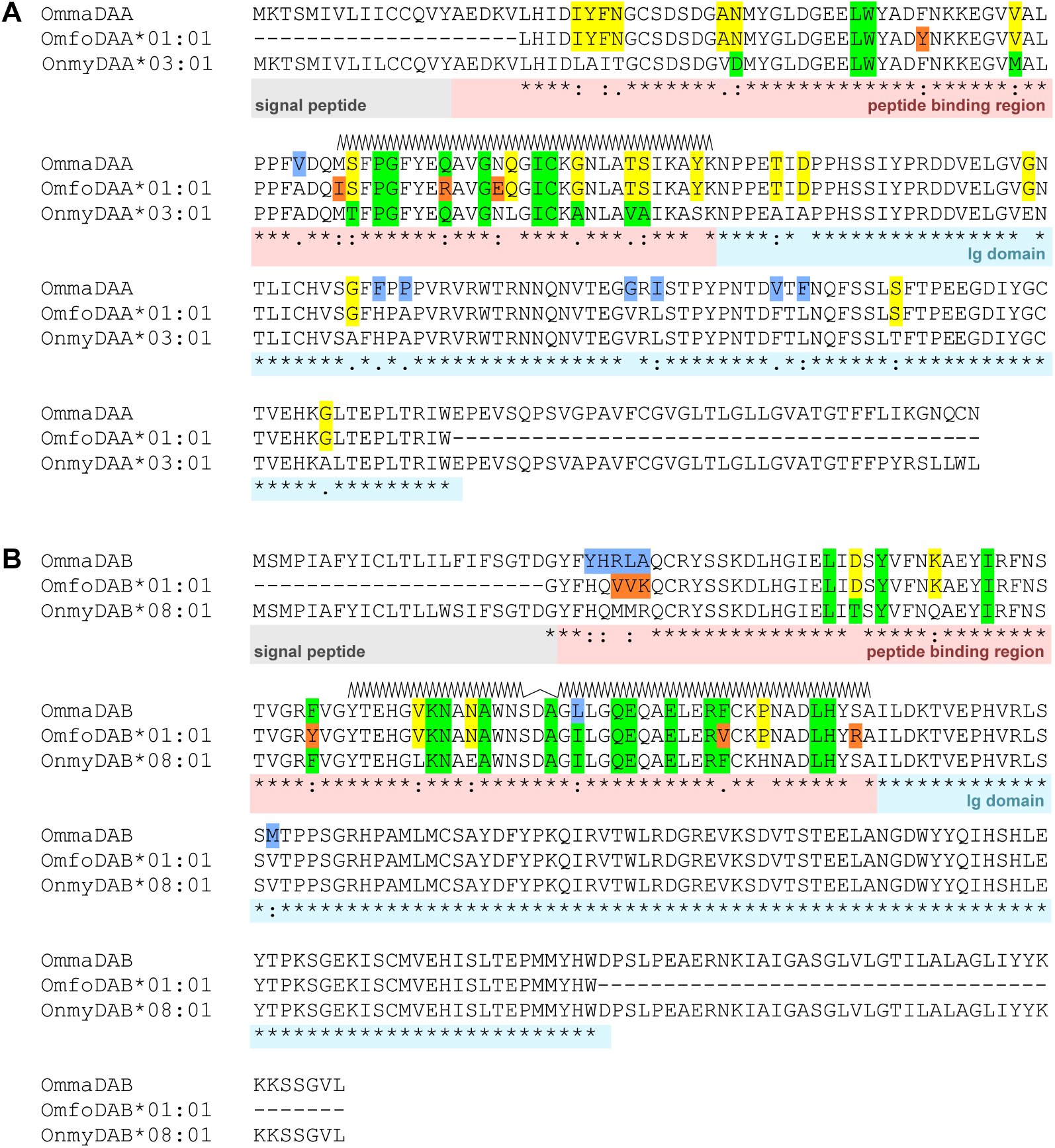
Alignments of the deduced classical MHC class II amino acid sequences. CLUSTALW was used to align the *O. m. formosanus* DAA/DAB alleles (determined in this study) and *O. m. masou* DAA(XP_064817619.1)/DAB(XP_064817620.1) to (**A**) OnmyDAA*03:01 (IPD-MHC_FISH08121) and (**B**) OnmyDAB*08:01 (IPD-MHC_FISH08133) of *O. mykiss*, respectively. The signal peptide, peptide binding region, and Ig domain are indicated below the alignments, and the alpha-helices forming the walls of the peptide binding groove based on an Alphafold structure prediction are shown above the sequences. Residues predicted to directly contact the antigen peptide binding based on (Gómez et al., 2010) are highlighted in green. Differences to the *O. mykiss* references unique to *O. m. masou* are highlighted in blue, unique to *O. m. formosanus* in orange, and common to both *O. masou* species in yellow, respectively.

Taken together, we cloned and identified only a single DAA (OmfoDAA*01:01) and DAB (OmfoDAB*01:01) allele from eight apparently unrelated individuals. The most parsimonious explanation is that they represent the dominant (if not exclusive) alleles for these loci and that additional variants (if they exist) occur at frequencies of less than 6.25% (1 out of 16). This lack of variation mirrors a recent microsatellite analysis in *O. m. formosanus* (Yamamoto et al., 2020) and our previous observation of a complete absence of genomic SNPs in six *O. m. formosanus* cytokine gene loci (Yen et al., 2024) that are autosomal in *O. mykiss.* Together this raises the concerning possibility that the remaining genetic diversity in the extant fish population is very limited.

### 3.2 The UBA gene in O. m.formosanus is polymorphic

To test whether the classical MHC class I gene (*UBA*) is also homozygous in *O. m. formosanus* we decided to clone the α1 region (encoded in exon 2) and α2 region (encoded in exon 3 and 4) that form the PBR of this MHC complex. To design PCR primer pairs to amplify exon 2 we selected regions upstream and downstream of this exon that are almost perfectly conserved between three Pacific salmon species (*O. mykiss*, *Oncorhynchus tshawytscha* [Chinook salmon], *Oncorhynchus gorbuscha* [pink salmon], and *O. m. masou*). As the sequence upstream of *UBA* exon 3, however, is poorly conserved and the complete genome assembly of *O. m. masou* had become available during the course of this study, we solely relied on its sequences for establishing the exon 3+4 primers. We again started with the genomic DNA of a single individual (FLS#775) as a template and this time were able to clone and sequence two distinct but highly similar (exon 2: 99%, exon 3+4: 97% identity) PCR products hereafter referred to as UBAα1*01:01, UBAα1*02:01, UBAα2*01:01, and UBAα2*02:01, respectively. As these sequences were highly similar to the *O. m. masou* genome (exon 2: 98-99%, exon 3+4: 97-98% identity, Fig. 3) it suggested that FLS#775 is heterozygous for the *UBA* gene and that at least two allelic variants exist in *O. formosanus*. While the similarities of the exon 2 genomic fragments to the *O. mykiss* (95%, LOC110496224), *O. tshawytscha* (92-93%, LOC112264791), and *O. gorbuscha* (93-94%, LOC124012358) *UBA* genes were in line with their evolutionary distance and our observations for DAA/DAB, the differences for the exon3+4 genomic fragments were considerably larger (Fig. 3). Only *O. gorbuscha* showed 91% identity over the entire length whereas the homologous regions detectable by BLAST was restricted to the two exonic regions for *O. mykiss* (87-89%) and only exon 4 for *O. tshawytscha* (90%). Yet these sequences were still the better matches within the respective genomes than any of the non-classical fish MHC class I genes including *UCA*, *UDA*, and *UGA*. Furthermore, it is important to consider that the *UBA* alleles present in the genome assemblies are random, and likely not the variants most closely related to those we cloned from *O. m. formosanus*.

**Figure 3.**
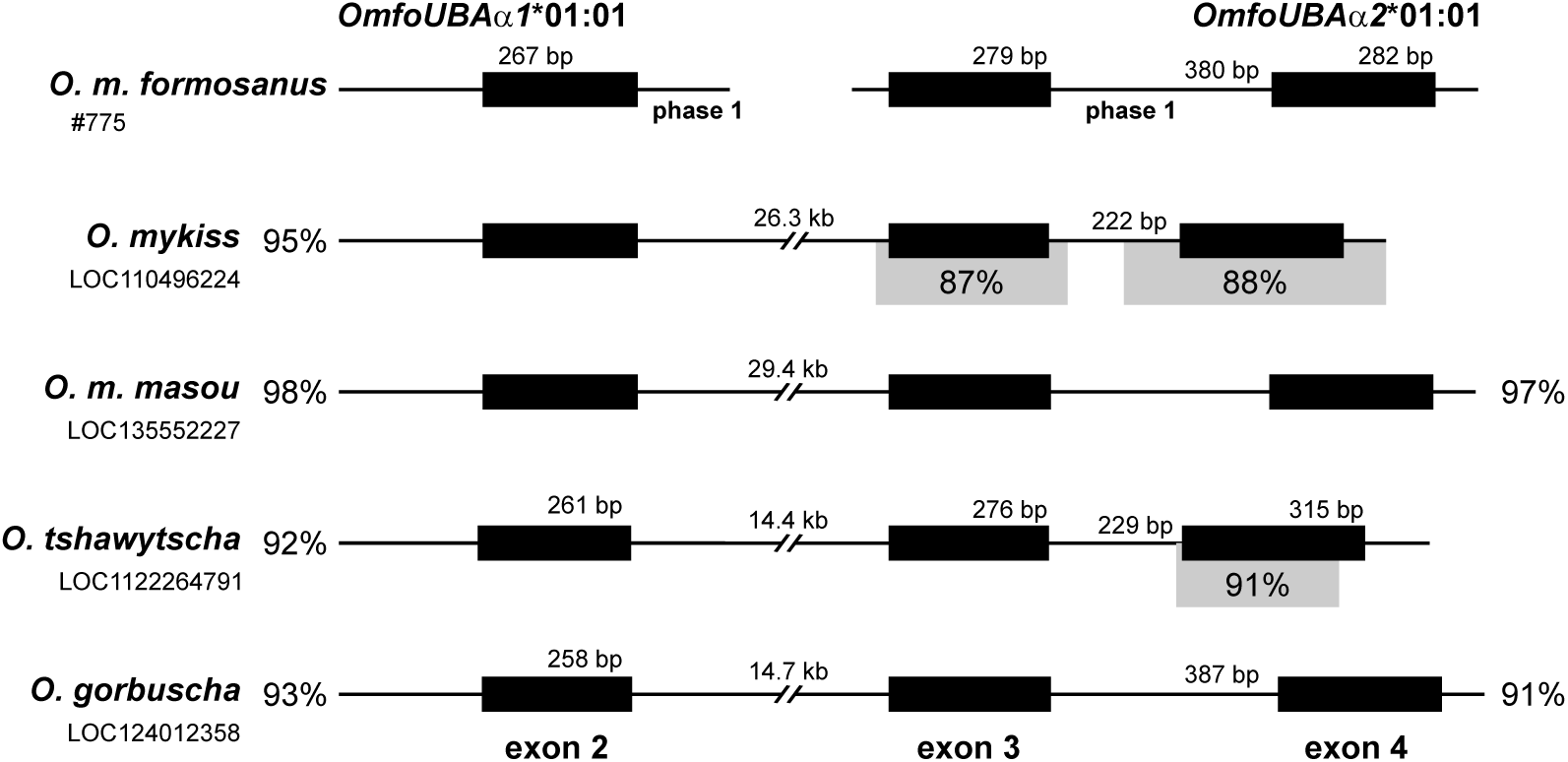
Comparison of the *UBA* gene loci of *O. m. formosanus* and related salmonid fish. The PCR amplicons of *OmfoUBA* exon 2 and exon 3+4 and the same regions in the current genome assemblies of *O. mykiss*, *O. m. masou*, *O. tshawytscha* (Otsh_v2.0, GCF_018296145.1), and *O. gorbuscha* (OgorEven_v1.0, GCF_021184085.1) are drawn to scale. The overall sequence identity to the *O. m. formosanus* PCR products (the most frequent allele is used here) compared to the sequences in the genome assemblies is shown in front (exon 2) or after (exon 3+4) each gene locus where applicable. In case of *O. mykiss* and *O. tshawytscha* only parts of the exon 3+4 region (marked in grey) show detectable sequence similarities. Exons are shown as filled boxes, the sizes of the exons and introns are shown, and the intron phases are noted below each intron. Note that all values identical to *O. m. formosanus* are being omitted for clarity.

To assess the allelic variation within *UBA* in our cohort, we cloned and sequenced exon 2 and exon 3+4 from each of the eight individuals. In contrast to the lack of diversity at the MHC class II loci, the nucleotide sequence alignment of both *UBA* PCR products from eight individual fish revealed a third allelic variant and resulting in overall allele frequencies of 62.5%, 31.25%, and 6.25%, respectively (Fig. 4). Notably all fish carried at least one copy of the most abundant allele for both amplicons (UBAα1*01:01 and UBAα2*01:01), with two fish (FLS#815 and FLS#842) appearing homozygous for this variant. The rarest set of alleles (UBAα1*03:01 and UBAα2*03:01), however, was only present in one single fish (FLS#810). As we were unable to amplify a larger PCR fragment harboring all three UBA exons (note that intron 2 is about 30 kb in length in the closely related *O. m. masou* genome), we do not have direct experimental data linking the sequence variants in exon 2 to those in exons 3+4. But given identical allele frequencies for both PCR products, and identical allele distributions for each fish, we inferred the linkage between the sequences of the two PCR products (Fig. 4). This allowed us to assemble the amino acid sequences of the entire peptide binding regions of three UBA variants (OmfoUBA*01:01, OmfoUBA*02:01, and OmfoUBA*03:01) encoded by our MHC class I amplicons. Importantly, the observed heterozygosities of (H_O_=0.750) for each PCR product and the assembled alleles, is much higher than the expected level H_E_=0.501 suggesting a selective advantage for heterozygosity. In summary, we discovered that the *OmfoUBA* gene locus retained important genetic diversity within the fish population despite the homozygosity of the classic MHC class II loci.

**Figure 4.**
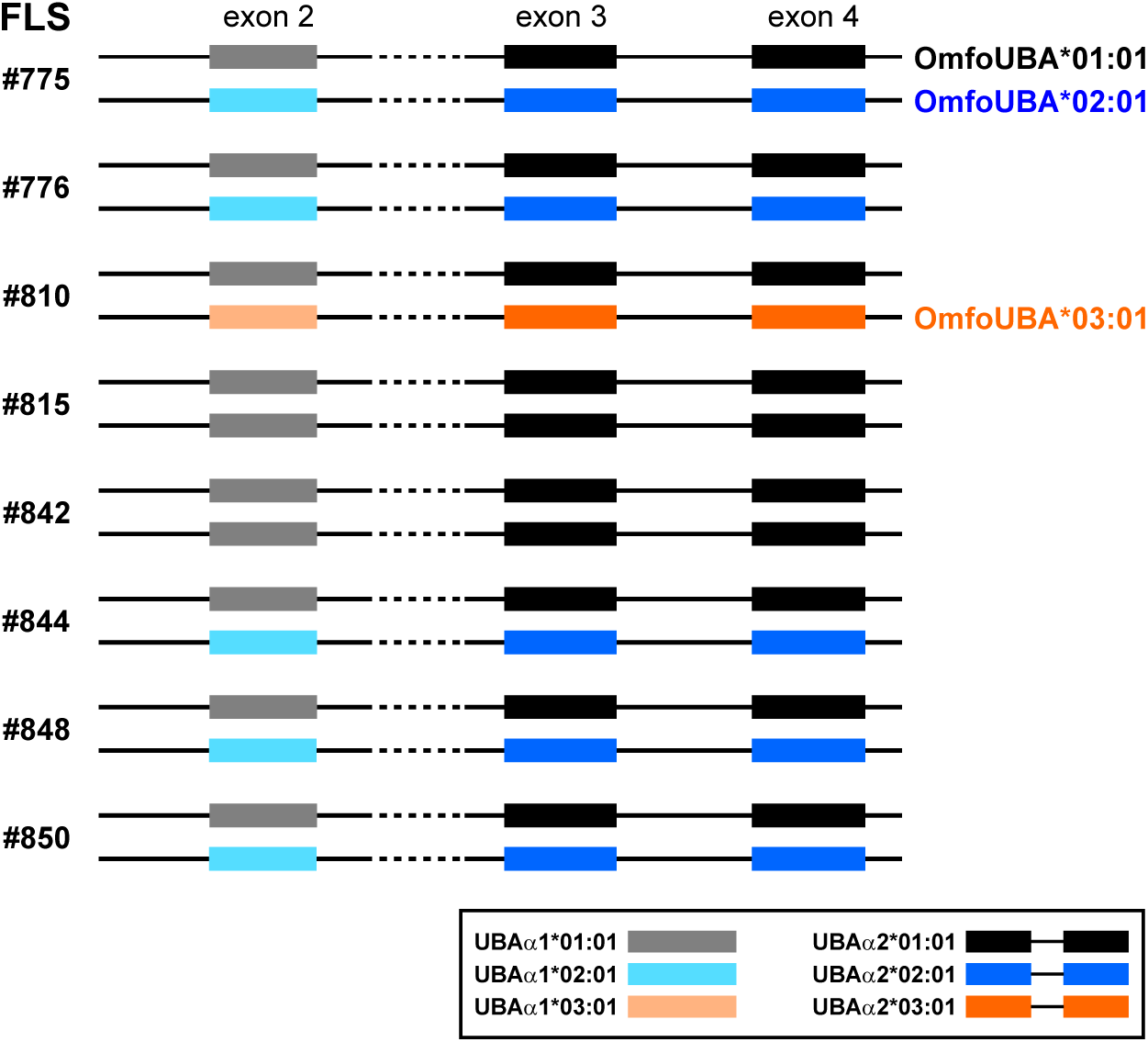
*UBA* allele distribution in our *O. m. formosanus* cohort. The genotypes for the exons 2 and exon 3+4 regions of the *OmfoUBA* gene are shown for each of eight individuals. As the allele distribution matches perfectly for both amplicons across all fish, we assembled them into three complete *OmfoUBA* alleles as shown on the right.

To extend our analysis to the protein level, we translated the coding sequences within each *OmfoUBA* allele into amino acid sequences, and a BLASTP search within the extensive MHC sequence datasets at GenBank revealed the highest similarities to the single *O. m. masou* UBA sequence (XP_064839799.1; OmfoUBA*01:01=93%, OmfoUBA*02:01=93%, OmfoUBA*03:01=95%). When looking separately at the α1 and α2 domains a wider range of sequences emerged as best hits for the former: UBAα1*01:01=91% with XP_064839799.1 (*O. m. masou*), UBAα1*02:01=90% with AAB62228.1 (*O. mykiss*), UBAα1*03:01=94% with CAK18621.1 (*Salmo trutta*, Satr-UBA*1801). In contrast all three OmfoUBAα2 alleles matched best to the UBAα2 of XP_064839799 (*O. m. masou*) with 91%, 93% and 95% identity, respectively. When comparing to the *O. mykiss* UBA alleles in the manually curated IPD-MHC database the closest homologs for OmfoUBA*01:01, OmfoUBA*02:01, and OmfoUBA*03:01 were OnmyUBA*13:01 (89%), OnmyUBA*11:01 (89%), and OnmyUBA*11:01 (91%), respectively. A similarly diverse set of best matches were observed when analyzing the α1 and α2 domains separately with five different *O. mykiss* UBA alleles (for α1: *01:03=84%, *01:03=90%, *16:01=93%; and for α2: *11:01=89%, *24:01=91%, *11:01=90%). Taken together this suggests that a diverse population of UBA variants exists in the extant *O. masou* populations (in particular in *O. m. formosanus*), and that these cannot be traced back to a single UBA allele in a common ancestor of all Pacific salmons as they likely evolved by recombination between multiple ancestral sequences.

To explore putative consequences of the UBA sequence variations in *O. m. formosanus* on peptide presentation, we performed a multiple sequence alignment that included the O. mykiss UBA*13:01 allele that is most similar to OmfoUBA*01:01 (Fig. 5). Overall, we observed differences at 30 positions with 25/30 in the peptide binding regions (14 in α1 and 11 in α2) and only 5/30 were observed in the Ig domain that resides at the base of the MHC class I extracellular domain. As expected, a significant number of the sequence variations locate to positions lining the antigen peptide binding groove with six in α1 and seven in α2, respectively. No differences between the three alleles were observed in the eight residues predicted to directly interact with the peptide ends, this was also the case when compared to the OnmyUBA*13:01 suggesting that these alleles share a common preference for the nature of the anchor residues of the peptides they present.

**Figure 5.**
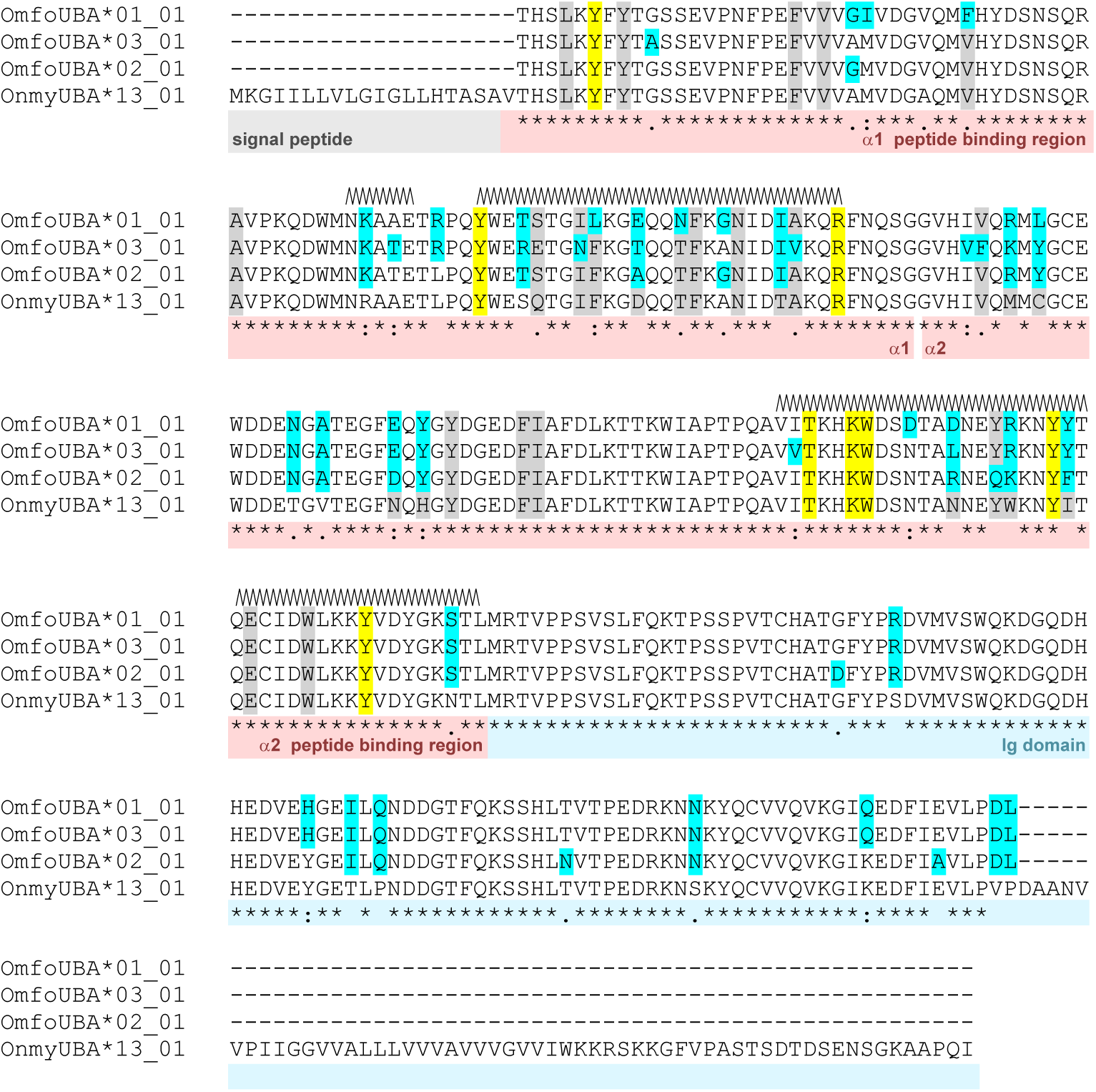
Alignments of the deduced classical MHC class I amino acid sequences. CLUSTALW was used to align the three assembled *O. m. formosanus* UBA alleles (determined in this study) to OnmyUBA*13:01 (IPD-MHC:FISH08166) of *O. mykiss*. respectively. The signal peptides, peptide binding regions, and Ig domain are indicated below the alignments, and the alpha-helical regions forming the walls of the antigen peptide binding groove are shown above the sequences. Residues forming this groove are marked in grey and conserved amino acids involved in contacting the peptide ends are highlighted in yellow following the nomenclature of (Yamaguchi and Dijkstra, 2019). Differences among the *O. m. formosanus* alleles and compared to *O. mykiss* are highlighted in cyan.

## 4. Discussion

Here we report first sequence data for the classical MHC genes *DAA*, *DAB*, and *UBA* genes in *O. m. formosanus*. The evolutionary history of this species includes several events that likely created genetic bottle necks that are expected to be reflected in reduced genetic diversity in the extant population. A recent report and our previous analysis of *O. m. formosanus* cytokine genes already indicated a surprisingly high level of homozygosity (Yamamoto et al., 2020; Yen et al., 2024), and in here it is even more apparent for the *OmfoDAA* and *OmfoDAB* genes that appear to be homozygous and identical in eight unrelated fish that were collected in the wild. Notably these sequences were highly similar to yet still different from the *DAA* and *DAB* genes in the genome assembly of *O. m. masou* from Japan (the closest relative to the Taiwanese salmon) suggesting that there is diversity within these genes among the different *O. masou* species across the Northeast Pacific. While the formal possibility remains that our primer pairs were only able to pick up a single allele for each gene, this is a rare scenario for two primer pair designed based on genomic sequences conserved across *Oncorhynchus* species from the East and West Pacific within genes that are otherwise highly polymorphic in salmonid fish. Alternatively, the DAA/DAB MHC complex could be non-classical (consistent with being monomorphic) in *O. m. formosanus* and we will screen for polymorphisms in DAA/DAB and additional MHC class II genes (like DBA/DBB) in a larger cohort and in Japanese *O. masou* in the future to address this question.

Reduced genetic diversity is in general considered a concern for the survival of a population and in one such instance, the mammoth on Wrangel island in Siberia, has been linked to the extinction of a species (Rogers and Slatkin, 2017). Even arthropods that propagate by parthenogenesis using unfertilized eggs that should result in a loss of heterozygosity, avoid the detrimental consequences by ensuring the co-inheritance of both parental alleles (Lacy et al., 2024). Beyond the broad negative effects of increased homozygosity, what are the specific functional consequences of a monomorphic DAA/DAB pair? As only a single type of MHC class II complex can be formed (in contrast to the four variants that are encoded by heterozygous *DAA* and *DAB* alleles) it clearly limits the range of antigen peptides that can be presented to CD4^+^ T helper cells by professional antigen presenting cells. This is thought to impair adaptive immune response to extracellular pathogens whose peptides cannot be presented by the single classical MHC class II variant. There is, however, no published evidence for immune defects within the *O. m. formosanus* population thus far.

In contrast to the loss of diversity in *DAA*/*DAB*, there are at least three alleles of the single classical MHC class I gene *UBA* in the extant population and we anticipate that future surveys of a larger cohort will reveal additional alleles. The dichotomy between monomorphic MHC class II and polymorphic class I in *O. m. formosanus* is enabled by the fact that unlike in mammals these genes are unlinked in teleost fish (Bingulac-Popovic et al., 1997). Among the three *OmfoUBA* alleles one is dominant such that it is present in every single individual we tested suggesting that it provides an important protection against a distinct set of viruses or intracellular bacteria in this species. Furthermore, the observed level of heterozygosity is higher than expected which is in line with the idea that the ability to present a larger set of antigen peptides provides a survival benefit (Pierini and Lenz, 2018). But although some QTLs for disease resistance map to *MHC* loci in *O. mykiss* (Ozaki et al., 2001), none appears to be linked to *UBA* gene (Yamaguchi and Dijkstra, 2019). There is, however, an unexpected genetic link between *O. mykiss* behaviour and this classical *MHC* class I gene locus (Azuma et al., 2005). Finally, a comprehensive study of the *UBA* alleles in a larger cohort of *O. m. formosanus* and the Japanses *O. masou* species will be required to gain insight into whether different UBA variants emerged in response to different environments that they now inhabit since their split after the last glacial period.

One important conclusion from our findings is that although genomic diversity appears limited it clearly still exists in the extant *O. m. formosanus* population. Notably we thus far only found genetic variations in one of the gene loci that is general considered to be the most polymorphic locus in vertebrates, a classical MHC class I gene (Jin et al., 2018), whereas the equally likely candidates for such diversity, the classical MHC class II genes, were monomorphic. We predict that there are likely other polymorphic loci within the *O. m. formosanus* genome and genes in the vicinity of *UBA* are among the candidates. A systematic genome-wide population genetics analysis will be required to address this question. Yet, the *UBA* is already one suitable marker to study the genetic diversity of *O. m. formosanus*. As our cohort was sampled in 2016 and since then nursery stock was introduced to successfully and rapidly increase the numbers of fish in the wild, it is warranted to begin assessing changes in genetic diversity of the population until now. Such approach could also be applied to the breeding program itself to preferentially use breeding pairs that would maximize MHC class I diversity in their offspring.

## Acknowledgements

We thank all members of the Fugmann and Yang labs for helpful discussions. This work was supported by grants of the Chang Gung Memorial Hospital [CMRPD1N0312] to S.D.F and [CMRPD1M0462] to S.Y.Y. as well as private funds from the Fugmann/Yang family.

## References

Apanius, V., Penn, D., Slev, P.R., Ruff, L.R., Potts, W.K., 1997. The nature of selection on the major histocompatibility complex. Crit Rev Immunol 17, 179–224. 10.1615/CRITREVIMMUNOL.V17.I2.40

Azuma, T., Dijkstra, J.M., Kiryu, I., Sekiguchi, T., Terada, Y., Asahina, K., Fischer, U., Ototake, M., 2005. Growth and behavioral traits in Donaldson rainbow trout (Oncorhynchus mykiss) cosegregate with classical major histocompatibility complex (MHC) class I genotype. Behav Genet 35, 463–478. 10.1007/S10519-004-0863-6

Bingulac-Popovic, J., Figueroa, F., Sato, A., Talbot, W.S., Johnson, S.L., Gates, M., Postlethwait, J.H., Klein, J., 1997. Mapping of mhc class I and class II regions to different linkage groups in the zebrafish, Danio rerio. Immunogenetics 46, 129–134. 10.1007/S002510050251

Christensen, K.A., Flores, A.M., Joshi, J., Shibata, K., Fujimoto, T., Koop, B.F., Devlin, R.H., 2025. Masu salmon species complex relationships and sex chromosomes revealed from analyses of the masu salmon (Oncorhynchus masou masou) genome assembly. G3 Genes|Genomes|Genetics 15, 278. 10.1093/G3JOURNAL/JKAE278

Gómez, D., Conejeros, P., Marshall, S.H., Consuegra, S., 2010. MHC evolution in three salmonid species: a comparison between class II alpha and beta genes. Immunogenetics 62, 531–542. 10.1007/S00251-010-0456-X

Grimholt, U., Lukacs, M., 2021. Fate of MHCII in salmonids following 4WGD. Immunogenetics 73, 79–91. 10.1007/S00251-020-01190-6

Gwo, J.C., Hsu, T.H., Lin, K.H., Chou, Y.C., 2008. Genetic relationship among four subspecies of cherry salmon (Oncorhynchus masou) inferred using AFLP. Mol Phylogenet Evol 48, 776–781. 10.1016/J.YMPEV.2007.12.023

Healey, M., Kline, P., Tsai, C.-F., 2011. Saving the Endangered Formosa Landlocked Salmon. Fisheries Magazine 26, 6–14.

Jin, Y., Wang, J., Bachtiar, M., Chong, S.S., Lee, C.G.L., 2018. Architecture of polymorphisms in the human genome reveals functionally important and positively selected variants in immune response and drug transporter genes. Hum Genomics 12. 10.1186/S40246-018-0175-1

Lacy, K.D., Hart, T., Kronauer, D.J.C., 2024. Co-inheritance of recombined chromatids maintains heterozygosity in a parthenogenetic ant. Nature Ecology & Evolution 2024 8:8 8, 1522–1533. 10.1038/s41559-024-02455-z

Larkin, M.A., Blackshields, G., Brown, N.P., Chenna, R., Mcgettigan, P.A., McWilliam, H., Valentin, F., Wallace, I.M., Wilm, A., Lopez, R., Thompson, J.D., Gibson, T.J., Higgins, D.G., 2007. Clustal W and Clustal X version 2.0. Bioinformatics 23, 2947–2948. 10.1093/BIOINFORMATICS/BTM404

McClelland, E.K., Ming, T.J., Tabata, A., Miller, K.M., 2011. Sequence analysis of MHC class I α2 from sockeye salmon (Oncorhynchus nerka). Fish Shellfish Immunol 31, 507–510. 10.1016/J.FSI.2011.06.012

Morita, K., 2011. Masu Salmon Group, in: Beamish, R.J. (Ed.), Ocean Ecology of Pacific Salmon and Trout. American Fisheries Society, Bethesda, USA.

Ozaki, A., Sakamoto, T., Khoo, S., Nakamura, K., Coimbra, M.R.M., Akutsu, T., Okamoto, N., 2001. Quantitative trait loci (QTLs) associated with resistance/susceptibility to infectious pancreatic necrosis virus (IPNV) in rainbow trout (Oncorhynchus mykiss). Mol Genet Genomics 265, 23–31. 10.1007/S004380000392

Pierini, F., Lenz, T.L., 2018. Divergent Allele Advantage at Human MHC Genes: Signatures of Past and Ongoing Selection. Mol Biol Evol 35, 2145–2158. 10.1093/MOLBEV/MSY116

Rogers, R.L., Slatkin, M., 2017. Excess of genomic defects in a woolly mammoth on Wrangel island. PLoS Genet 13, e1006601. 10.1371/JOURNAL.PGEN.1006601

Sommer, S., 2005. The importance of immune gene variability (MHC) in evolutionary ecology and conservation. Front Zool 2, 16. 10.1186/1742-9994-2-16

Stern, L.J., Wiley, D.C., 1994. Antigenic peptide binding by class I and class II histocompatibility proteins. Structure 2, 245–251. 10.1016/S0969-2126(00)00026-5/ASSET/C8023C60-0BDF-4AAF-84F3-7EBB5ACA1027/MAIN.ASSETS/GR6.JPG

Yamaguchi, T., Dijkstra, J.M., 2019. Major Histocompatibility Complex (MHC) Genes and Disease Resistance in Fish. Cells 8. 10.3390/CELLS8040378

Yamamoto, S., Morita, K., Kikko, T., Kawamura, K., Sato, S., Gwo, J.C., 2020. Phylogeography of a salmonid fish, masu salmon Oncorhynchus masou subspecies-complex, with disjunct distributions across the temperate northern Pacific. Freshw Biol 65, 698–715. 10.1111/FWB.13460

Yen, Y.H., Zheng, D.Y., Yang, S.Y., Gwo, J.C., Fugmann, S.D., 2024. The cytokine genes of Oncorhynchus masou formosanus include a defective interleukin-4/13A gene. Dev Comp Immunol 155. 10.1016/J.DCI.2024.105156

