## Supplementary Table 1 for "The classical MHC class I and II genes of *O. m. formosanus* exhibit different polymorphism levels"

### Primers for cloning *O. m. formosanus* MHC gene loci

| Target | Primer | Sequence | Product size |
|---|---|---|---|
| *UBA exon 2* | O-UBAe2f | ACATGGTGTCTTAGCAACTTTC | 672 bp |
| | O-UBAe2r | TCTGTACATCCCTGATATTCT | |
| *UBA exon 3+4* | UBA-NF1 | GCCAGAAGAATGCACAGAGT | 1073 bp |
| | UBA-NR2 | TAGAAATTAAGAATATTGGGCAC | |
| *DAA exon 2+3* | DAA-F | GTTCATGTTGGTAAAGCAGGACAG | 894 bp |
| | DAA-R | AAGAAGTCAAAAGTAATGCAGGCT | |
| *DAB exon 2+3* | DAB-F | TTGGCAGAGTACCACATTTAACA | 1218 bp |
| | DAB-R | TATCGCCCCTTACCCCAGTG | |

**Supplementary Table 1**
