## Supplementary Data File 1 for "The classical MHC class I and II genes of *O. m. formosanus* exhibit different polymorphism levels"

**Amino acid sequences of the DAA, DAB and UBA alleles of O. m. formosanus**

```
>OmfoDAA*01:01
LHIDIYFNGCSDSDGANMYGLDGEELWYADYNKKEGVVALPPFADQISFP
GFYERAVGEQGICKGNLATSIKAYKNPPETIDPPHSSIYPRDDVELGVGN
TLICHVSGFHPAPVRVRWTRNNQNVTEGVRLSTPYPNTDFTLNQFSSLSF
TPEEGDIYGCTVEHKGLTEPLTRIW


>OmfoDAB*01:01
GYFHQVVKQCRYSSKDLHGIELIDSYVFNKAEYIRFNSTVGRYVGYTEHG
VKNANAWNSDAGILGQEQAELERVCKPNADLHYRAILDKTVEPHVRLSSV
TPPSGRHPAMLMCSAYDFYPKQIRVTWLRDGREVKSDVTSTEELANGDWY
YQIHSHLEYTPKSGEKISCMVEHISLTEPMMYHW


>OmfoUBAα1*01:01
THSLKYFYTGSSEVPNFPEFVVVGIVDGVQMFHYDSNSQRAVPKQDWMNK
AAETRPQYWETSTGILKGEQQNFKGNIDIAKQRFNQSG

>OmfoUBAα1*02:01
THSLKYFYTGSSEVPNFPEFVVVGMVDGVQMVHYDSNSQRAVPKQDWMNK
ATETLPQYWETSTGIFKGAQQTFKGNIDIAKQRFNQSG

>OmfoUBAα1*03:01
THSLKYFYTASSEVPNFPEFVVVAMVDGVQMVHYDSNSQRAVPKQDWMNK
ATETRPQYWERETGNFKGTQQTFKANIDIVKQRFNQSG


>OmfoUBAα2*01:01
GVHIVQRMLGCEWDDENGATEGFEQYGYDGEDFIAFDLKTTKWIAPTPQA
VITKHKWDSDTADNEYRKNYYTQECIDWLKKYVDYGKSTL

>OmfoUBAα2*02:01
GVHIVQRMYGCEWDDENGATEGFDQYGYDGEDFIAFDLKTTKWIAPTPQA
VITKHKWDSNTARNEQKKNYFTQECIDWLKKYVDYGKSTL

>OmfoUBAα2*03:01
GVHVFQKMYGCEWDDENGATEGFEQYGYDGEDFIAFDLKTTKWIAPTPQA
VVTKHKWDSNTALNEYRKNYYTQECIDWLKKYVDYGKSTL


>OmfoUBA*01:01 assembled_exon2+3+4
THSLKYFYTGSSEVPNFPEFVVVGIVDGVQMFHYDSNSQRAVPKQDWMNK
AAETRPQYWETSTGILKGEQQNFKGNIDIAKQRFNQSGGVHIVQRMLGCE
WDDENGATEGFEQYGYDGEDFIAFDLKTTKWIAPTPQAVITKHKWDSDTA
DNEYRKNYYTQECIDWLKKYVDYGKSTLMRTVPPSVSLFQKTPSSPVTCH
ATGFYPRDVMVSWQKDGQDHHEDVEHGEILQNDDGTFQKSSHLTVTPEDR
KNNKYQCVVQVKGIQEDFIEVLPDL

>UBA*02:01 assembled_exon2+3+4
THSLKYFYTGSSEVPNFPEFVVVGMVDGVQMVHYDSNSQRAVPKQDWMNK
ATETLPQYWETSTGIFKGAQQTFKGNIDIAKQRFNQSGGVHIVQRMYGCE
WDDENGATEGFDQYGYDGEDFIAFDLKTTKWIAPTPQAVITKHKWDSNTA
RNEQKKNYFTQECIDWLKKYVDYGKSTLMRTVPPSVSLFQKTPSSPVTCH
ATDFYPRDVMVSWQKDGQDHHEDVEYGEILQNDDGTFQKSSHLNVTPEDR
KNNKYQCVVQVKGIKEDFIAVLPDL

>UBA*03:01 assembled_exon2+3+4
THSLKYFYTASSEVPNFPEFVVVAMVDGVQMVHYDSNSQRAVPKQDWMNK
ATETRPQYWERETGNFKGTQQTFKANIDIVKQRFNQSGGVHVFQKMYGCE
WDDENGATEGFEQYGYDGEDFIAFDLKTTKWIAPTPQAVVTKHKWDSNTA
LNEYRKNYYTQECIDWLKKYVDYGKSTLMRTVPPSVSLFQKTPSSPVTCH
ATGFYPRDVMVSWQKDGQDHHEDVEHGEILQNDDGTFQKSSHLTVTPEDR
KNNKYQCVVQVKGIQEDFIEVLPDL
```
